# Robust diagnosis of Ewing sarcoma by immunohistochemical detection of super-enhancer-driven EWSR1-ETS targets

**DOI:** 10.1101/158766

**Authors:** Michaela C. Baldauf, Martin F. Orth, Marlene Dallmayer, Aruna Marchetto, Julia S. Gerke, Rebeca Alba Rubio, Merve M. Kiran, Julian Musa, Maximilian M. L. Knott, Shunya Ohmura, Jing Li, Nusret Akpolat, Ayse N. Akatli, Özlem Özen, Uta Dirksen, Wolfgang Hartmann, Enrique de Alava, Daniel Baumhoer, Giuseppina Sannino, Thomas Kirchner, Thomas G. P. Grünewald

## Abstract

Ewing sarcoma is an undifferentiated bone-associated cancer. Although molecular detection of pathognomonic *EWSR1-ETS* fusions such as *EWSR1-FLI1* enables definitive diagnosis, substantial confusion can arise if molecular diagnostics are unavailable. Diagnosis based solely on the conventional immunohistochemical marker CD99 is unreliable due to its abundant expression in morphological mimics. This study aimed to identify novel diagnostic immunohistochemical markers for Ewing sarcoma.

We analyzed 768 expression microarrays representing 21 tumor entities including Ewing-like sarcomas to nominate candidate biomarkers. These candidates were validated by immunohistochemistry (IHC) in a tissue microarray (TMA) comprising 174 samples. Microarray, chromatin immunoprecipitation and sequencing (ChIP-Seq) data, and reporter assays were employed to analyze their EWSR1-FLI1-dependency.

Our comparative expression analyses revealed that *ATP1A1, BCL11B*, and *GLG1* constitute specific markers for Ewing sarcoma. Analysis of ChIP-Seq and microarray datasets showed that their expression is EWSR1-FLI1-dependent. This outcome corresponded to EWSR1-FLI1-binding to proximal super-enhancers, which showed high activity in reporter assays. Consistently, high ATP1A1, BCL11B, and GLG1 expressions were detected by IHC. Automated cut-off-finding and combination-testing in the TMA demonstrated that detection of high BCL11B and/or GLG1 expression is sufficient to reach 96% specificity for Ewing sarcoma. While 88% of tested Ewing-like sarcomas displayed strong CD99-immunoreactivity, none displayed combined high expression of BCL11B and GLG1.

Collectively, we provide evidence that *ATP1A1, BCL11B*, and *GLG1* are EWSR1-FLI1 targets, of which BCL11B and GLG1 offer a fast, simple and cost-efficient way to diagnose Ewing sarcoma by IHC. We anticipate that these markers will significantly reduce the number of misdiagnosed patients, and thus improve patient care.

## INTRODUCTION

Ewing sarcoma is characterized by the presence of chimeric *EWSR1-ETS* fusion oncogenes ^1^. Before the discovery of this unifying genetic hallmark, diagnosing Ewing sarcoma definitively was challenging ^2^ as Ewing sarcoma tumors are largely composed of undifferentiated cells displaying a small-blue-round cell phenotype ^3,4^ This phenotype is shared by many other tumor entities such as rhabdomyosarcoma and neuroblastoma ^5^. Recently, several so-called Ewing-like sarcoma sub-types have been identified ^6–9^. These tumors are characterized by distinct fusion oncogenes and transcriptomic signatures ^6–12^, as well as (most likely) by distinct clinical behavior ^6,12,13^.

Although Ewing sarcoma can usually be reliably distinguished from its morphological mimics by cytogenetic and molecular genetic analyses ^14,15^, there is currently no robust biomarker available for routine histology. Substantial diagnostic confusion can arise because sophisticated cytogenetic and molecular diagnostic techniques are not universally available or too expensive for some diagnostic facilities (particularly in developing countries). While the only established immunohistochemical biomarker for Ewing sarcoma, CD99, shows high sensitivity, its low specificity and high expression in morphological mimics such as CIC-and BCOR-rearranged sarcomas, as well as in certain lymphoma subtypes and poorly differentiated synovial sarcoma, are problematic ^3,11–13,16–18^. Thus, CD99 alone is unreliable to definitively diagnose Ewing sarcoma.

In the current study, comparative expression analyses revealed that *ATP1A1, BCL11B,* and *GLG1* constitute potential specific markers for Ewing sarcoma. Expression of these genes appeared to be induced by EWSR1-FLI1-bound super-enhancers, which showed high activity in reporter assays. Specific immunohistochemical staining of these proteins in comprehensive tissue microarrays (TMAs), combined with automated cut-off determination and combination-testing, demonstrated that detecting high BCL11B and/or GLG1 levels is sufficient to reach 96% specificity for Ewing sarcoma. In fact, these markers were extremely effective at discriminating Ewing sarcoma from Ewing-like sarcomas.

Hence, these results provide a fast, simple and cost-efficient means of diagnosing Ewing sarcoma by immunohistochemistry (IHC), which is a considerable advantage for diagnostic facilities where molecular diagnostics are not available. This finding may significantly reduce the number of misdiagnosed patients and thus improve patient care.

## MATERIALS AND METHODS

### Human samples and ethics approval

Human tissue samples were retrieved from the archives of the Institute of Pathology of the LMU Munich (Germany), the Department of Pathology, Turgut Ozal Medical Center, Inonu University (Turkey), the Baçkent University Hospital (Turkey), the Gerhard-Domagk-Institute for Pathology of the University of Münster (Germany), the Institute of Biomedicine of Seville (Spain), and the Bone Tumour Reference Centre at the Institute of Pathology of the University Hospital Basel (Switzerland) with approval of the corresponding institutional review boards. The LMU Munich’s ethics committee approved the current study (approval no. 550-16 UE).

### Microarray analyses

Publicly available gene expression data generated with the Affymetrix HG-U133Plus2.0 DNA microarray for 1,790 samples comprising 21 tumor entities and 71 normal tissue types were retrieved from several repositories. Accession codes are given in **Suppl. Table 1**. All Ewing sarcoma samples were genetically verified to contain a specific *EWSR1-ETS* translocation as previously described ^19^. After rigorous quality-checks (including the Relative Log Expression (RLE) and Normalized Unscaled Standard Error (NUSE)) and careful clinical annotation validation, expression intensities were calculated simultaneously with the Robust Multi-array Average (RMA) algorithm (including background adjustment, quantile normalization and summarization, using custom brainarray CDF (ENTREZG, v19)), which yielded one optimized probe-set per gene ^20^. The pairwise expression ratio (ER) of every gene was calculated based on its median expression levels in primary Ewing sarcoma tumors and any of the 20 other remaining tumor entities. The differential gene expression’s statistical significance was calculated with an unpaired, two-tailed Student’s t-test. The resulting *P* values were adjusted for multiple testing with the Bonferroni method. Only genes with an ER of > 2 between Ewing sarcoma and any other tumor entity and a Bonferroni-corrected *P* value < 0.05 across all tumor entities compared with Ewing sarcoma were considered diagnostically relevant.

Publicly available gene expression microarray data for ectopic EWSR1-FLI1 expression in embryonic stem cells (Affymetrix HG-U133Plus2.0; GSE64686 ^21^) and from Ewing sarcoma cell lines that were either transiently transfected with an shRNA directed against EWSR1-FLI1 or a control shRNA (TC252, SK-N-MC, STA-ET-7.2, STA-ET-1, WE68; Affymetrix HG-U133A; GSE14543 ^22^) or stably transduced with a doxycycline-inducible shRNA against EWSR1-FLI1 (A673; Affymetrix HG-U133A 2.0; GSE27524 ^23^) were normalized by RMA using custom brainarray CDF (ENTREZG, v19).

To identify the pathways and biological processes associated with a given gene present in normalized gene expression data from primary Ewing sarcoma tumors, gene-set enrichment analyses (GSEAs) were performed on ranked lists of genes in which all genes were ranked by their correlation coefficient with the given reference gene (MSigDB, c2.all.v5.1). GSEA was carried out with 1,000 permutations in default settings ^24^.

**Analysis of DNase-Seq and chromatin immunoprecipitation followed by high-throughput sequencing (ChIP-Seq) data and genome-wide identification of superenhancers**

Publicly available data were retrieved from the Gene Expression Omnibus (GEO). ENCODE SK-N-MC DNase-Seq (GSM736570) ^25^ were analyzed in the Nebula environment ^26^ using Model-based Analysis of ChIP-Seq v1.4.2 (MACS) ^27^ and converted to *.wig format for display in the UCSC Genome Browser ^28^. Preprocessed ChIP-Seq data from Riggi *et al.* ^29^ (GSE61944) were converted to *.wig format with the UCSC’s bigWigToWig conversion tool. The following samples were used in this study:

- ENCODE_SKNMC_hg19_DNAseHS_rep2
- GSM1517546 SKNMC.shGFP96.FLI1
- GSM1517555 SKNMC.shFLI196.FLI1
- GSM1517547 SKNMC.shGFP96.H3K27ac
- GSM1517556 SKNMC.shFLI196.H3K27ac
- GSM1517569 A673.shGFP48.FLI1
- GSM1517572 A673.shFLI148.FLI1
- GSM1517571 A673.shGFP96.H3.k27ac
- GSM1517574 A673.shFLI196.H3K27ac

ChIP-seq data of the histone modification H3K27ac in A673 and SK-N-MC Ewing sarcoma cell lines (shGFP96) from a genome-wide chromatin analysis (GSE61944) conducted by Riggi *et al.* ^29^ was used for epigenetic analysis of enhancers. The already aligned Sequence Read Archives (*.sra) of both cell lines and the corresponding whole cell extracts were downloaded from GEO. Before peak calling with MACS2 ^27^, the data were prepared with SAMtools ^30^. ChIP peak annotation was done with HOMER ^31^. The super-enhancers were identified with ROSE ^32,33^.

### Cell culture, DNA constructs, and reporter assays

A673/TR/shEF1 Ewing sarcoma cells, which harbor a doxycycline-inducible shRNA against EWSR1-FLI1, were described previously ^34^ and kindly provided by J. Alonso (Madrid, Spain). A673 cells were obtained from ATCC. All cells were grown at 37°C in 5% CO_2_ in a humidified atmosphere in RPMI 1640 medium (Biochrom) containing 10% Tetracycline-free FCS (Biochrom), 100 U/ml penicillin, and 100 p,g/ml streptomycin (both Biochrom). Cell line purity was confirmed by short tandem repeat profiling (latest profiling 15^th^ December 2015), and cells were checked routinely for the absence of mycoplasma by PCR. Human GGAA-microsatellites close to the *ATP1A1, BCL11B,* or *GLG1* gene were cloned from the A673 Ewing sarcoma cell line into the pGL3-luc vector (Promega) upstream of the SV40 minimal promoter. qPCR-Primer sequences were as follows:

- forward 5’-CTAGCCCGGGCTCGAGAGCAACACAAGGACTCAATTAC-3’ and reverse 5’-GATCGCAGATCTCGAGCTACTATGATGCAAAGCTGAGTG-3’ for the *ATP1A1* associated GGAA-microsatellite;
- forward 5’-CTAGCCCGGGCTCGAGGCCGTCTCTCTGTTCCTTAT-3’ and reverse 5’-GATCGCAGATCTCGAGAATCTCTGCTCCTTCATCCC-3’ for the *BCL11B* associated GGAA-microsatellite; and
- forward 5’-CTAGCCCGGGCTCGAGGCTACTATAGCCAAATGCAAAGAAGAA-3’ and reverse 5’-GATCGCAGATCTCGAGTGCACTGGGTTATACAGAAAGAGTTC-3’ for the *GLG1* associated GGAA-microsatellite.

For the reporter assays, 3 × 10^5^ A673/TR/shEF1 cells per well of a six-well plate were seeded in 2.5 ml medium and transfected with pGL3-luc vectors and *Renilla* pGL3-Rluc (ratio, 100:1) using Lipofectamine LTX and Plus Reagent (Invitrogen). After 4 h transfection media were replaced by media with or without doxycycline (1 p,g/ml). Cells were lysed after 72 h and assayed with a dual luciferase assay system (Berthold). *Firefly* luciferase activity was normalized to *Renilla* luciferase activity.

### RNA extraction, reverse transcription, and quantitative real-time PCR (qRT-PCR)

RNA was extracted with the Nucleospin II kit (Macherey-Nagel) and reverse-transcribed using the High-Capacity cDNA Reverse Transcription Kit (Applied Biosystems). qPCRs were performed using SYBR green (Applied Biosystems). Oligonucleotides were purchased from MWG Eurofins Genomics. Reactions were run on a Bio-Rad CFX Connect instrument and analyzed using Bio-Rad CFX Manager 3.1 software.

### Construction TMAs and IHC

A total of 174 archival formalin-fixed and paraffin-embedded (FFPE) primary tissue samples with reviewed histological diagnosis were obtained from the participating institutions and collected at LMU Munich’s Institute of Pathology. Representative FFPE tumor blocks were also selected for TMA construction at LMU Munich’s Institute of Pathology. A detailed description of the TMA is given in **Suppl. Table 2**.

All Ewing sarcoma FFPE samples showed cytogenetic evidence for a translocation of the *EWSR1* gene as determined by fluorescence in situ hybridization (FISH) and were reviewed by a reference pathologist. For this study, Ewing-like sarcomas were defined as small-blue-round cell sarcomas being either positive for *CIC-DUX4* (8 cases) or *BCOR-CCNB3* (2 cases) or unclassified (7 cases) after extensive reference pathologist work-up. Each TMA slide contained three cores (each 1 mm in diameter) from every sample as well as internal controls.

For IHC, 4 µm sections were cut, and antigen retrieval was performed with microwave treatment with 750W at pH7.5 TRIS buffer (2 × 15 min) using the antigen retrieval AR kit (DCS, HK057-5KE) for GLG1 or the Target Retrieval Solution (Dako, S1699) for BCL11B and ATP1A1. Blockage of endogenous peroxidase was performed using 7.5% aqueous H_2_O_2_ solution at room temperature and blocking serum from the corresponding kits for 20 min.

Slides were then incubated for 60 min with the primary antibodies anti-ATP1A1 (1:330 dilution, Proteintech, 14418-1-AP) ^35^, anti-BCL11B (1:1000 dilution, Abcam, ab70453), or anti-GLG1 (1:250 dilution, Sigma, HPA010815) ^36^. Then slides were incubated with a secondary anti-rabbit IgG antibody (ImmPress Reagent Kit, Peroxidase-conjugated) followed by target detection using AECplus chromogen for 10 min (Dako, K3461).

For IHC of CD99, 4-p.m sections were cut and incubated for 32 min with the antibody anti-CD99 (1:40 dilution, Dako, 12E7) using the Roche UltraView detection kit.

### Evaluation of immunoreactivity and automated cut-off finding

Semi-quantitative evaluation of marker immunostaining was carried out by three independent observers (MCB, MD, MFO) analogous to scoring of hormone receptor Immune Reactive Score ranging from 0-12 according to Remmele and Stegner ^37^, which is routinely used in surgical pathology to quantify hormone receptor expression in mammary carcinoma.

The percentage of cells with high expression was scored and classified in five grades (grade 0 = 0–19%, grade 1 = 20–39%, grade 2 = 40–59%, grade 3 = 60–79% and grade 4 = 80–100%). The fraction of cells showing high expression was scored after examination of 10 high-power fields (40x) of at least one section per sample. In addition, the intensity of marker immunoreactivity was determined (grade 0 = none, grade 1 = low, grade 2 = moderate and grade 3 = strong). The product of these two grades defined the final immunoreactivity score. Sensitivity and specificity of each marker for Ewing sarcoma were calculated with an inhouse generated VBA (Visual Basic for Applications) script implemented in Microsoft Excel (Microsoft). The script computed sensitivity and specificity for all possible combinations of markers and within these combinations, for all possible cut-offs for every marker. The best marker and cut-off combination was chosen based on the following criteria: high specificity (defined as > 95%), high sensitivity, and discriminability between positive (IRS higher than the cut-off) and negative samples.

### Survival analysis

Microarray data of 166 primary Ewing sarcoma tumors (GSE63157 ^38^, GSE34620 ^19^, GSE12102 ^39^, and GSE17618 ^40^), which had well-curated clinical annotations available, were downloaded from the GEO. The data were generated on Affymetrix HG-U133Plus2.0 or Affymetrix HuEx-1.0-st microarray chips and normalized separately by RMA using custom brainarray CDF files (v20). Batch effects were removed using ComBat ^41,42^. Samples were stratified into two groups based on their median intra-tumoral gene expression levels. Significance levels were calculated with a Mantel-Haenszel test. *P* values < 0.05 were considered statistically significant.

## RESULTS

### *ATP1A1, BCL11B,* and *GLG1* are strongly overexpressed in Ewing sarcoma compared to tumor entities of differential diagnostic relevance

To identify highly specific diagnostic markers for Ewing sarcoma, publicly available microarray gene expression data comprising genetically confirmed Ewing sarcoma samples^19^, 20 additional tumor entities of potential differential diagnostic relevance ^5^, and 71 normal tissue types we retrieved. Based on these microarray expression data the median expression of every gene represented on the Affymetrix HG-U133Plus2 microarray was determined. Next, we calculated the expression ratio (ER) for every gene based on its median expression in pairwise comparisons of Ewing sarcoma and the remaining tumor entities. Only genes, which were strongly overexpressed in Ewing sarcoma compared to all other tumor entities defined by a minimal log2-transformed ER of >2, were considered as diagnostically relevant. Of the 19,702 genes represented on the microarray platform, 51 had an ER of > 2 across all tested tumor entities. In parallel, the level of significance of the differential expression of all genes in pairwise comparisons of Ewing sarcoma relative to all other tumor entities was calculated. Only 10 genes exhibited a Bonferroni-corrected *P* value < 0.05 (**Fig. 1a**, **b**). Next, both gene lists were crossed, which showed that only 3 genes, termed *ATP1A1* (ATP1A1 Na^+^/K^+^ transporting subunit alpha), *BCL11B* (B-cell CLL/lymphoma 11B), and *GLG1* (Golgi glycoprotein 1) were both strongly and highly significantly overexpressed in Ewing sarcoma compared to all other tumor entities (**Fig. 1b**).

**Figure 1.**
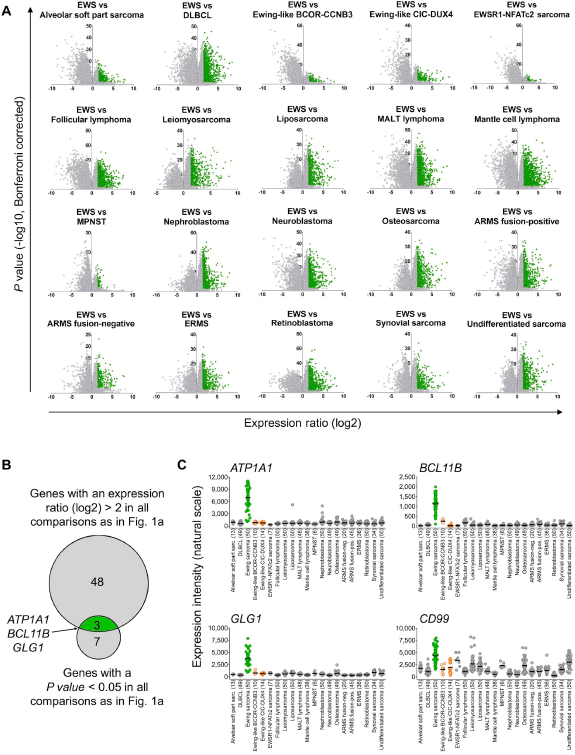
***ATP1A1, BCL11B,* and *GLG1* are strongly overexpressed in Ewing sarcoma compared to tumor entities of differential diagnostic relevance** A) Volcano plots of pairwise comparison of gene expression in Ewing sarcoma (EWS) and indicated tumor entities: diffuse large B-cell lymphoma (DLBCL); mucosa-associated lymphoid tissue (MALT) lymphoma; malignant peripheral nerve sheath tumor (MPNST); alveolar rhabdomyosarcoma (ARMS); and embryonal rhabdomyosarcoma (ERMS). Genes represented in green color had an expression ratio > 2 (log2) and a *P* value < 0.05 (Bonferroni-corrected). B) Size-proportional Venn diagram showing the overlap of genes differentially and significantly (minimal log2 expression ratio >2; *P* value < 0.05, Bonferroni corrected) overexpressed in Ewing sarcoma relative to all other tumor entities given in A and C. C) Scatter dot plot depicting gene expression levels of *ATP1A1, BCL11B, GLG1,* and *CD99* as determined by Affymetrix HG-U133Plus2.0 microarrays in primary tumors of 21 different entities. Ewing sarcoma is highlighted in green, Ewing-like sarcomas *(CIC-DUX4* or *BCOR-CCNB3* translocation positive) are highlighted in orange. Horizontal bars represent median expression levels. The number of analyzed samples is given in parentheses.

Then, the expression profiles of these three candidate biomarkers were compared to the conventional Ewing sarcoma marker *CD99* across all tumor entities. While *CD99* showed broad expression in many different tumor entities, *ATP1A1, BCL11B,* and *GLG1* were only expressed at low levels in every tumor entity relative to Ewing sarcoma, indicating a higher specificity for this disease than *CD99* (**Fig. 1c**).

Because a mixture of tumor tissue and normal cells, which could express the three markers could complicate immunohistochemical evaluation, we explored the expression levels of *ATP1A1, BCL11B,* and *GLG1* and that of *CD99* in Ewing sarcoma samples relative to 71 normal tissue types comprising 998 samples. As displayed in **Suppl. Fig. 1**, *ATP1A1, BCL11B,* and *GLG1* were only lowly expressed in some normal tissue types, while *CD99* was rather broadly expressed across many normal tissue types.

### EWSR1-FLI1 induces *ATP1A1, BCL11B,* and *GLG1* expression by binding to GGAA-microsatellites found in super-enhancers

The specific expression of the three candidate biomarkers in primary Ewing sarcoma suggests a possible regulatory relationship between them and EWSR1-FLI1. In fact, *ATP1A1* and *GLG1* were previously shown to be upregulated after ectopic expression of *EWSR1-FLI1* in the rhabdomyosarcoma cell line RD ^43^. Moreover, *BCL11B* was shown to be upregulated by EWSR1-FLI1 in Ewing sarcoma cell lines ^44^.

To further explore this regulatory relationship, available gene expression data was assessed, which showed that the ectopic EWSR1-FLI1 expression in embryonic stem cells was sufficient to significantly induce the expression of *ATP1A1, BCL11B,* and *GLG1* (**Fig. 2a**). Conversely, the shRNA-mediated knockdown of EWSR1-FLI1 in six different Ewing sarcoma cell lines significantly decreased their expression levels (**Fig. 2b**). Such consistent EWSR1-FLI1-dependent regulation was not observed for *CD99* (**Fig. 2a**, **b**).

**Figure 2.**
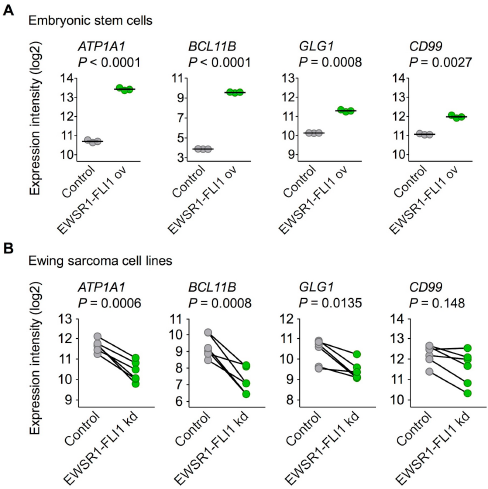
**EWSR1-FLI1 is sufficient to induce *ATP1A1, BCL11B,* and *GLG1* expression** A) Analysis of gene expression levels of *ATP1A1, BCL11B, GLG1* and *CD99* by Affymetrix HG-U133Plus2.0 microarrays in human embryonic stem cells after ectopic expression of EWSR1-FLI1 (GSE64686). Bars represent the medians. Two-tailed student’s T test. B) Analysis of gene expression levels of *ATP1A1, BCL11B, GLG1* and *CD99* by Affymetrix HG-U133A microarrays 96 h after short hairpin RNA-mediated knockdown of EWSR1-FLI1 in six different Ewing sarcoma cell lines (GSE14543 and GSE27524). Data are represented as before-after plots in which each dot represents a cell line. Two-tailed student’s T test.

These data in cell lines suggested that *ATP1A1, BCL11B,* and *GLG1* may be direct EWSR1-FLI1 target genes. Testing this hypothesis involved analyzing available ChIP-Seq and DNase-Seq data generated in Ewing sarcoma cell lines, which showed strong EWSR1-FLI1-binding to GGAA-microsatellites close to these genes. Notably, these GGAA-microsatellites exhibit characteristics of active EWSR1-FLI1-dependent enhancers (**Fig. 3a**). In fact, EWSR1-FLI1 is known to convert non-functional GGAA-microsatellites into potent enhancers to steer a large proportion of its target genes ^45–47^ Strong EWSR1-FLI1-dependent enhancer activity of these GGAA-microsatellites in luciferase reporter assays was consistently observed (**Fig. 3b**). In agreement with previous observations ^48^, these EWSR1-FLI1-dependent enhancers showed the typical H3K27ac profile of so-called super-enhancers in the A673 and SK-N-MC Ewing sarcoma cell lines (**Fig. 3c**, **Suppl. Tables 3 & 4**), which are considered to have tissue-defining functions ^32,49^. In addition to these findings *in vitro,* gene-set enrichment analyses of either *ATP1A1-, BCL11B-,* or GLG1-correlated genes within 166 primary Ewing sarcoma tumors revealed that the most significantly (min. NES = 3.08, *P* < 0. 001, q < 0.001) associated gene expression signature among the 3,687 tested was for each candidate marker ‘ZHANG_TARGETS_OF_EWSR1-FLI1_FUSION’ ^43^ (**Suppl. Table 5**).

**Figure 3.**
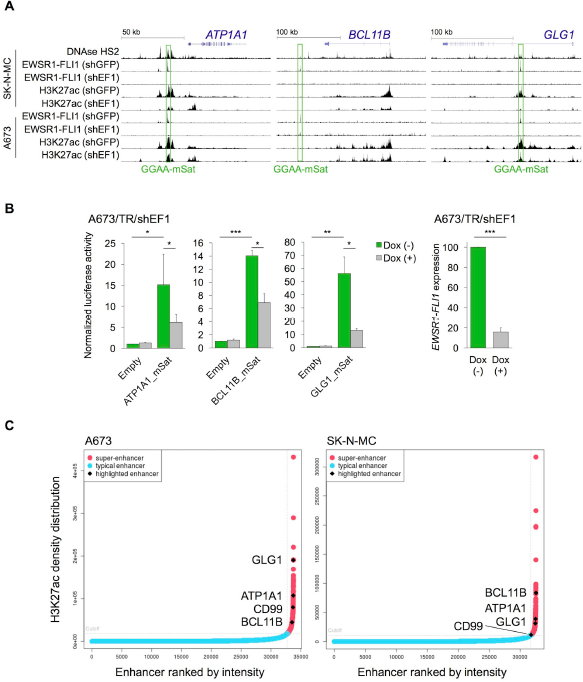
**EWSR1-FLI1 binds to GGAA-microsatellites with enhancer activity located close to or within the *ATP1A1, BCL11B,* or *GLG1* gene** A) Published DNase-Seq and ChIP-Seq data generated in Ewing sarcoma cell lines were displayed in the UCSC genome browser. shGFP, control; shEF1, shEWSR1-FLI1. GGAA-mSat, GGAA-microsatellite. B) Luciferase reporter assays in A673/TR/shEF1 cells containing a doxycycline (dox)-inducible shRNA against EWSR1-FLI1 confirmed the EWSR1-FLI1-dependent enhancer activity of cloned GGAA-microsatellites (1 kb fragments). EWSR1-FLI1 knockdown was confirmed by qRT-PCR 72 h after transfection. Data are presented as mean and SEM of *n* = 3 independent experiments. Two-tailed student’s t-test. * *P* < 0.05; ** *P* < 0.01; *** *P* < 0.001. C) Genome-wide analysis of published H3K27ac profiles of A673 and SK-N-MC Ewing sarcoma cell lines (GSE61944) identified super-enhancers proximal to *ATP1A1, BCL11B, GLG1,* and *CD99.* Enhancers are ranked by their H3K27ac density.

Collectively, these data strongly suggest that *ATP1A1, BCL11B,* and *GLG1* are direct EWSR1-FLI1 target genes.

### *ATP1A1* and *GLG1* may have prognostic relevance in Ewing sarcoma

To explore the potential of *ATP1A1, BCL11B, GLG1,* and *CD99* as prognostic biomarkers, we analyzed the association of their expression levels with outcome in a large sample of Ewing sarcoma patients (*n* = 166). Whereas higher *ATP1A1* and *GLG1* expression levels showed a significant correlation with better patient outcome (*P* = 0.006 and *P* = 0.0028, respectively), *BCL11B* and *CD99* did not (Suppl. Fig. 2).

### High expression of BCL11B and/or GLG1 is sufficient to robustly diagnose Ewing sarcoma by IHC

To confirm the overexpression of ATP1A1, BCL11B, and GLG1 on the protein level a comprehensive TMA including many solid tumor entities closely resembling Ewing sarcoma and other sarcoma entities was generated (**Suppl. Table 2**). Immunohistochemical staining of the TMAs was carried out with anti-ATP1A1, anti-BCL11B, anti-GLG1 and anti-CD99 antibodies, and immunoreactivity scores (IRS) were determined using the Remmele and Stegner ^37^ scoring system (IRS range from 0 to 12; **Fig. 4a**, **b**). As displayed in **Fig. 4b**, CD99 expression was not very specific for Ewing sarcoma compared to other sarcoma entities as well as Ewing-like sarcomas. However, CD99 reached 100% sensitivity for Ewing sarcoma in this TMA when applying a cut-off of IRS > 2. Compared to CD99, the three candidate markers were all less sensitive at any given cut-off, but much more specific (specificity 90 – 97%) when being highly expressed (defined as IRS > 9).

**Figure 4.**
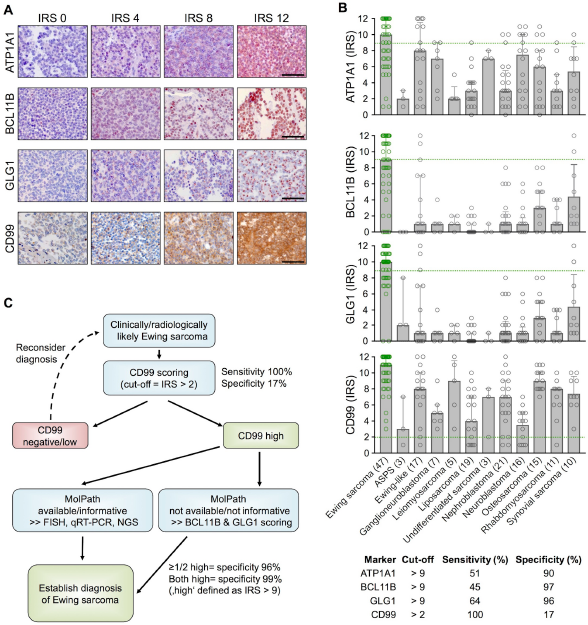
**High expression of BCL11B and/or GLG1 is sufficient to robustly diagnose Ewing sarcoma by IHC** A) Representative IHC images of Ewing sarcoma samples for the indicated marker. ATP1A1 is expressed in the cytoplasm, BCL11B in the nucleus, GLG1 at the perinuclear Golgi apparatus, and CD99 at the membrane. Scale bars = 100 μm. B) Scatter dot plots of the individual IRS for the indicated marker. The number of analyzed samples is given in parentheses. Bars represent mean IRS values, whiskers indicate the 95%-CI. Green dashed lines indicate the cut-offs to define sensitivity as specificity detecting for Ewing sarcoma as given in the table below. Alveolar soft part sarcoma (ASPS). C) Proposed work-flow for establishing robust diagnosis of Ewing sarcoma.

Automated cut-off-finding and combination-testing algorithms were then applied to the samples and set of candidate markers to identify a minimal set of markers and optimal cutoffs for robustly diagnosing Ewing sarcoma by IHC. These analyses indicated that while CD99 is a very valuable marker for screening for Ewing sarcoma, it needs auxiliary markers to establish a robust diagnosis. Further analyses indicated that while ATP1A1 exhibited high specificity (90%), it had no additional value for establishing Ewing sarcoma diagnosis if it was combined with BCL11B and GLG1. In fact, detecting high BCL11B and/or GLG1 expression in CD99-high tumors reached a specificity for Ewing sarcoma of at least 96%, and of 99% if both markers were highly expressed (defined as a IRS > 9). Strikingly, strong combined immunoreactivity for BCL11B and GLG1 was not observed in any of the tested Ewing-like sarcomas, while CD99 was found in 15 of 17 cases (88%).

Thus, the following work-flow is proposed to establish a diagnosis of Ewing sarcoma (**Fig. 4c**): In the case of clinically and/or radiologically suspected Ewing sarcoma, a biopsy should first be stained for CD99. If CD99 is positive (defined as IRS > 2), confirmatory molecular diagnostic procedures (such as FISH, qRT-PCR, and/or next-generation sequencing), if available, are preferred. If molecular diagnostic procedures are unavailable or the biopsy material is not suitable, an IHC-staining for BCL11B and GLG1 as well as subsequent scoring according to the Remmele and Stegner system should be performed. Since high expression of BCL11B and/or GLG1 (defined as IRS > 9) was found in 79% of all Ewing Sarcoma cases and associated with a specificity of 96%, diagnosis of Ewing sarcoma should be strongly considered if one or both markers are highly expressed.

Collectively, our data provide evidence that fast and robust diagnosis of Ewing sarcoma is enabled by immunohistochemical detection of the super-enhancer-driven EWSR1-ETS targets BCL11B and GLG1.

## DISCUSSION

Ewing sarcoma is genetically defined by pathognomonic *EWSR1-ETS* fusion transcripts ^1^. To date, at least 18 types of chimeric *EWSR1-FLI1* transcripts have been reported ^6^. Alternatively, *EWSR1* can be fused with *ERG, ETV1, E1A-F* or *FEV* in Ewing sarcoma^6^.

Although several molecular diagnostic tools are available to identify Ewing sarcoma among morphological mimics by detecting these gene fusions (e.g. by FISH, qRT-PCR, and/or direct sequencing), there are several limitations: All these techniques require good-quality DNA or RNA, which is not available in more than 10% of cases ^11^. In addition, FISH can sometimes yield non-informative results ^14^ Moreover, there is a risk of false positives because break-apart of the *EWSR1* gene can also be observed in other sarcoma entities such as desmoplastic small round cell tumor, clear cell sarcoma, angiomatoid fibrous histiocytoma, extraskeletal myxoid chondrosarcoma, and a subset of myxoid liposarcoma ^50^. Conversely, PCR-based assays can yield false negative results as the PCR may not cover the entire spectrum of different *EWSR1-ETS* fusions. Thus, some authors recommend combining FISH and qRT-PCR ^11^. However, these sophisticated techniques are not available in all diagnostic facilities, especially in developing countries, which poses a significant obstacle accurately diagnosing Ewing sarcoma.

To offer a simple, fast, and cost-effective way to reliably diagnose Ewing sarcoma by IHC, we combined *in silico*, *in vitro*, and *in situ* analyses, and found that the high expression of BCL11B and/or GLG1 is nearly diagnostic for this disease. It was shown that both genes are direct EWSR1-FLI1-targets, which are specifically overexpressed in Ewing sarcoma. In fact, their genetic loci exhibit EWSR1-FLI1-dependent super-enhancers that usually control the expression of tissue-defining genes ^32^. In particular, the high expression of the chosen markers was highly effective in discriminating Ewing sarcoma from EWSR1-ETS-negative Ewing-like sarcomas, which expressed *CD99* at high levels in 88% of our cases. Previously, another EWSR1-FLI1 target gene, *NKX2-2,* was proposed to serve in combination with CD99 as a useful immunohistochemical marker for Ewing sarcoma ^51^. In our comparative microarray analyses, *NKX2-2* did not, however, meet the stringent selection criteria for further validation. Similarly, another report showed that NKX2-2 is not fully specific for Ewing sarcoma ^52^.

Although most Ewing sarcoma tumors show little infiltration by lymphocytes ^53^, the fact that *BCL11B* is expressed in normal T cells (**Suppl. Fig. 1**) should be taken into account when assessing immunoreactivity in small-round-cell tumors that *BCL11B* is expressed in normal T cells. In indeterminate cases, a CD3 staining may be helpful.

Interestingly, all three original candidate markers play a role in fibroblast growth factor (FGF)-signaling. ATP1A1 is required for unconventional secretion of FGF ^54^, BCL11B promotes FGF-signaling by transcriptional suppression of a negative feedback inhibitor ^44,55^, and GLG1 (alias cysteine-rich FGF receptor) is known to regulate intracellular levels of FGF^56^. Several studies have shown that FGF promotes EWSR1-FLI1 expression ^57^ and growth of Ewing sarcoma cells *in vitro* and *in vivo* ^46,55^, and that FGF-inhibitors could be used as a targeted treatment for Ewing sarcoma patients ^58^. Although more work needs to be done to elucidate the precise role of ATP1A1, BCL11B, and GLG1 in FGF-signaling, it is tempting to speculate that they could serve as predictive biomarkers for the efficacy of FGF-inhibitors.

Collectively, we propose utilizing BCL11B and GLG1 as novel biomarkers for the diagnosis of Ewing sarcoma and recommend validating their diagnostic value in a prospective and multi-centered setting. It will be essential to further develop and characterize specific monoclonal antibodies directed against these proteins to improve and standardize their diagnostic utility.

## ACKNOWLEDGEMENTS

We would like to thank Mrs. Andrea Sendelhofert, Mrs. Anja Heier, and Mrs. Mona Melz for excellent technical assistance. The authors would like to thank all the donors and the Hospital Universitario Virgen del Rocío-Instituto de Biomedicina de Sevilla Biobank (Andalusian Public Health System Biobank and ISCIII-Biobank Platform PT13/0010/0056) for the human specimens used in this study. This work was supported by the German National Academic Foundation (to M.C.B.), the Kind-Philipp-Foundation (to J.M., M.C.B., M.F.O., M.D., A.M, and G.S.), the ‘Deutsche Stiftung für Junge Erwachsene mit Krebs’ (to M.D.), the German National Academic Foundation (to M.C.B. and MMLK), the ‘Verein zur Förderung von Wissenschaft und Forschung an der Medizinischen Fakultät der LMU München’ (WiFoMed; to T.G.P.G.), the Daimler and Benz Foundation in cooperation with the Reinhard Frank Foundation (to T.G.P.G.), the Friedrich-Baur-Stiftung (to T.G.P.G.) by LMU Munich’s Institutional Strategy LMUexcellent within the framework of the German Excellence Initiative (to T.G.P.G.), the ‘Mehr LEBEN für krebskranke Kinder – Bettina-Bräu-Stiftung’ (to T.G.P.G.), the Fritz-Thyssen Foundation (FTF-40.15.0.030MN, to T.G.P.G. and G.S.), the Wilhelm-Sander-Foundation (2016.167.1 to T.G.P.G.), and the German Cancer Aid (DKH-111886 and DKH-70112257 to T.G.P.G.; DKH-108128 to U.D.), EU FP7 and TRANSCAN EraNet - PROVABES [01KT1310], EU-FP7 EEC [602856-2] to U.D..

### AUTHORS’ CONTRIBUTIONS

MCB, MFO, MD, and TGPG conceived the study, wrote this paper, and drafted the figures and tables. MCB, JSG, MFO, and TGPG performed bioinformatic and statistical analyses. MMK, NA, ANA, 00, DB, EdA, UD, WH, and TK provided FFPE samples. MCB, MD, and MFO scored the TMAs. MMLK, RAR, SO, and JL helped in experimental procedures. TK provided laboratory infra-structure. AM, RAR, and GS cloned the GGAA-microsatellites and performed reporter assays. All authors read and approved the final manuscript.

### DISCLOSURE/DUALITY OF INTEREST

The authors declare no duality of interest.

## SUPPLEMENTARY FIGURE LEGENDS

**Figure 1.**
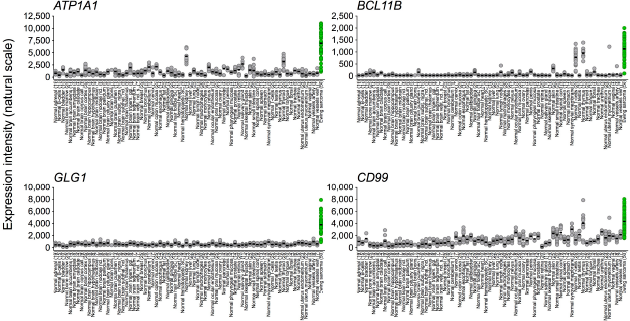
**Comparison of *ATP1A1, BCL11B,* and *GLG1* mRNA expression in Ewing sarcoma and normal tissues** Microarray data were normalized simultaneously by RMA using custom brainarray CDF files (v19) yielding one optimized probe-set per gene ^20^. Accession codes are given in Supplementary Table 1. Ewing sarcoma is highlighted in green color. Numbers of analyzed samples are given in parentheses.

**Figure 2.**
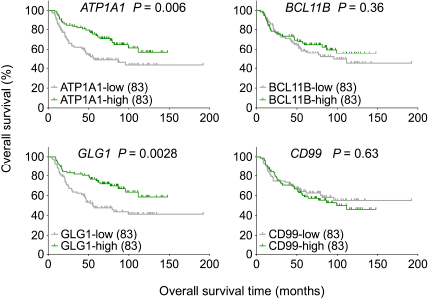
***ATP1A1* and *GLG1* may serve as prognostic biomarkers in Ewing sarcoma** Microarray data of 166 primary Ewing sarcoma tumors were normalized by RMA using custom brainarray CDF files (v20). Samples were stratified into two groups based on their median intratumoral gene expression levels. Significance levels were calculated with a log-rank test.

